# *xcore*: an R package for inference of gene expression regulators

**DOI:** 10.1101/2022.02.23.481130

**Authors:** Maciej Migdał, Cecilia Lanny Winata, Takahiro Arakawa, Satoshi Takizawa, Masaaki Furuno, Harukazu Suzuki, Erik Arner, Bogumił Kaczkowski

## Abstract

Elucidating the Transcription Factors (TFs) that drive the gene expression changes in a given experiment is one of the most common questions asked by researchers. The existing methods rely on the predicted Transcription Factor Binding Site (TFBS) to model the changes in the motif activity. Such methods only work for TFs that have a motif and assume the TF binding profile is the same in all cell types. Given the wealth of the ChIP-seq data available for a wide range of the TFs in various cell types, we propose that the modeling can be done using ChiP-seq “signatures” directly, effectively skipping the motif finding and TFBS prediction steps. We present *xcore*, an R package that allows Transcription Factor activity modeling based on their ChiP-seq signatures and user’s gene expression data. We also provide *xcoredata* a companion data package that provides a collection of preprocessed ChiP-seq signatures. We demonstrate that *xcore* leads to biologically relevant predictions using TGF-beta induced epithelial-mesenchymal transition and rinderpest infection time-course CAGE data as examples.

**Availability and Implementation:** *xcore* and *xcoredata* R packages are freely available at www.github.com/bkaczkowski/xcore and www.github.com/mcjmigdal/xcoredata.

*xcore* user guide is available www.bkaczkowski.github.io/xcore/articles/xcore_vignette.html.

## 1. Introduction

Gene expression profiling is often performed to elucidate the transcriptional regulators in a given system/perturbation. A common approach is to use transcription factor motifs to computationally predict the TFBS within promoter regions of known genes. The “motif activity” is then predicted/inferred based on gene expression profiles. (Balwierz *et al*., 2014; Schmidt *et al*., 2017; Madsen *et al*., 2018). Although such methods are quite simplistic, they proved useful for the identification of key molecular regulators (FANTOM Consortium *et al*., 2009; Natarajan *et al*., 2012; Balwierz *et al*., 2014; Schmidt and Schulz, 2019). The limitations are that many TFs do not have a defined motif and some binding events may be specific to a particular biological context.

ReMap (Chèneby *et al*., 2020) and ChIP-Atlas (Oki *et al*., 2018) provide a wealth of uniformly processed ChIP-seq data (genome-wide peaks) for TFs but also other transcriptional regulators including transcriptional coactivators and chromatin-remodeling factors. However, to our knowledge, there are no published methods that take advantage of this data to directly model the activity of transcriptional regulators.

Here, we propose to use the publicly available ChIP-seq data to directly represent the genome-wide occupancy of regulators. We intersected the peaks with promoter regions and used the linear ridge regression to infer the regulators associated with observed gene expression changes (Fig. 1A). The advantage of this approach is the direct integration of gene expression profiles with experimental TF binding data. We provide a) processed and pre-computed, ChIP-seq based molecular signatures (*xcoredata*), b) methodology for activity modeling (*xcore*). The framework is implemented as an R package (submitted to Bioconductor) and integrates smoothly with commonly used differential expression workflows like edgeR (Robinson, McCarthy and Smyth, 2010) or DESeq2 (Love, Huber and Anders, 2014).

**Figure 1.**
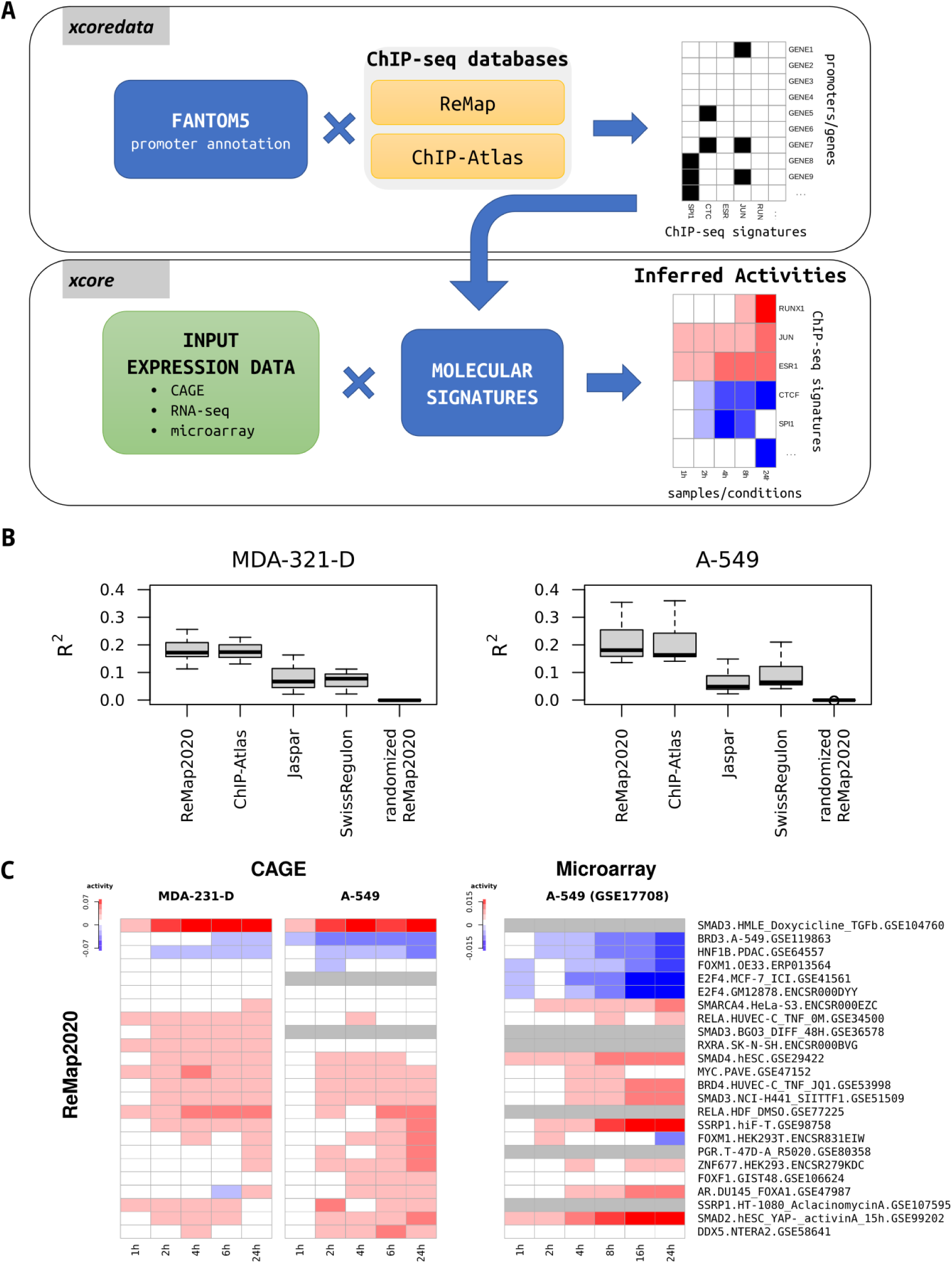
Inferring transcription factors activities from gene expression during TGFβ induced EMT in A-549 and MDA-231-D cell lines. **(A)** Flowchart depicts *xcore* and *xcoredata* functionalities. **(B)** Comparison of models describing gene expression changes between 0 and 24 hours after TGFβ treatment. Models were constructed using different molecular signature sets: Motif based (Jaspar, SwissRegulon) and ChIP-seq based (ReMap2020, ChIP-Atlas). **(C)** Heatmap showing the changes of TF activities versus 0h time point. The top-scoring ReMap2020 signatures are shown. Gray color designates NA values.

### 2. Materials and methods

### 2.1 Expression data processing

*xcore* takes promoter or gene expression counts matrix as input, the data is then filtered for lowly expressed features, normalized for the library size and transformed into counts per million (CPM) using edgeR (Robinson, McCarthy and Smyth, 2010). Users need to designate the base level samples by providing an experiment design matrix. These samples are used as a baseline expression when modeling changes in gene expression. CAGE data is an input of choice for promoter level analysis. However, *xcore* can be used with other types of expression data such as microarray or RNA-seq data to perform gene level analysis. Transcript level analysis based on RNA-seq data is possible in principle but currently not implemented.

### 2.2 Molecular signatures

A second input consists of molecular signatures describing known transcription factors binding preferences within the promoter’s vicinity. We provide sets of precomputed molecular signatures with *xcoredata*, the accompanying data package. The signatures were obtained by downloading all ChiP-seq data from ReMap2020 (Chèneby *et al*., 2020) and ChIP-Atlas (Oki *et al*., 2018) and intersecting it against +/- 500nt window of know promoter regions, defined based on FANTOM5’s hg38 annotation (Lizio *et al*., 2015, p. 5). The signatures can also be easily constructed using *xcore* by providing predicted TFBS or custom ChiP-seq peaks (see *xcore* user guide).

### 2.3 Expression modeling

Using ridge regression (Hoerl and Kennard, 1970) *xcore* models changes in expression as a linear combination of molecular signatures in an attempt to find their unknown activities. In layman’s terms, we are seeking to find a combination of ChiP-seq based signatures that best explains the changes of gene expression observed in a given experiment. Next, their estimated activities can be tested for significance (Cule, Vineis and De Iorio, 2011). This allows the selection of molecular signatures with the highest predicted influence on the expression changes.

To compare different models, coefficients of determination (R^2^) were calculated for selected time points using 10-fold cross-validation (CV) and pooling estimates across replicates.

## 3. Results

We used *xcore* to perform gene expression modeling analysis in the context of two CAGE datasets: TGF induced epithelial-mesenchymal transition (EMT) experiment performed in A-549 and MDA-231-D cell lines, FANTOM5’s rinderpest infection series dataset (Lizio *et* al., 2015, p. 5) and a microarray dataset: TGF induced EMT in A-549 cell line (GSE17708) (Sartor *et al*., 2010). First, we compared the models based on ChIP-seq signatures (ReMap2020 and ChIP-Atlas) vs motif based signatures (Jaspar and SwissRegulon). We observed that models based on ChiP-seq signatures showed on average higher R^2^ values, which reflects the proportion of variance explained by the model and overall “goodness of fit”. (Fig. 1B, Supp. Fig. 1B). To investigate the biological relevance of the obtained results, we looked at top-scoring signatures from ReMap2020 (Fig. 1C) and ChIP-Atlas (Supp. Fig, 1A) in TGF induced EMT datasets. Among those, we identified known key TFs involved in the TGF pathway such as *SMAD2/3/4* (Xu, Lamouille and Derynck, 2009), *SSRP1, HNF1B* (Lavin and Tiwari, 2020), *DDX5* (Dardenne *et al*., 2014, p. 5) *or RELA* (Tian *et al*., 2018). Other well-known EMT-linked TFs also returned as significant included *ZEB1, SNAI2, TBX3, SOX4* (Supp. Table 1, 2, 3). In case of FANTOM5’s rinderpest infection dataset, top-scoring ReMap2020 and ChIP-Atlas signatures (Supp. Fig. 2) showed several TFs involved in the closely related measles infection pathway, including *RELA, IRF9, TP53*, and *STAT1* (KEGG PATHWAY:map05162) (Kanehisa *et al*., 2010).

## 4. Discussion

*xcore* provides a flexible framework for integrative analysis of gene expression and publicly available TF binding data to unravel putative transcriptional regulators and their activities. Our analyses showed the superior results when using ChIP-seq based signatures as compared to motifs based ones. We attribute this difference to the presence of biotype specific binding information which might be lost in motifs that describe more general transcription factor binding preferences. Moreover, despite high numbers of ChIP-seq signatures and redundancy our machine learning framework is able to select biologically relevant signatures. In conclusion we believe that *xcore* is useful for hypothesis generation data and will be of use for many researchers.

## Acknowledgements

We thank Dr. Iga Jancewicz for insightful comments on the manuscript. We thank Dr. Daizo Koinuma and Dr. Kohei Miyazono (Department of Molecular Pathology, Graduate School of Medicine, The University of Tokyo, Japan) for their kindly providing MDA-231-D cells.

## Funding

This work was supported by a research grant from the Ministry of Education, Culture, Sport, Science and Technology of Japan for the RIKEN Center for Integrative Medical Sciences. MM was supported by RIKEN’s IMS Internship Program. MM is recipient of the Postgraduate School of Molecular Medicine doctoral fellowship financed by the European Union through the European Regional Development Fund under Knowledge Education Development programme within the project “Next generation sequencing technologies in biomedicine and personalized medicine”.

## Contribution

BK and EA conceived the study. TA, ST, MF and HS generated the data on TGF induced EMT in A-549 and MDA-231-D cell lines. MM and BK contributed to the design of the study and analyzed the data. MM wrote the R package. MM, CLW, EA and BK wrote the manuscript. All authors have read and approved the manuscript.

---

**Supplementary Figure 1.**
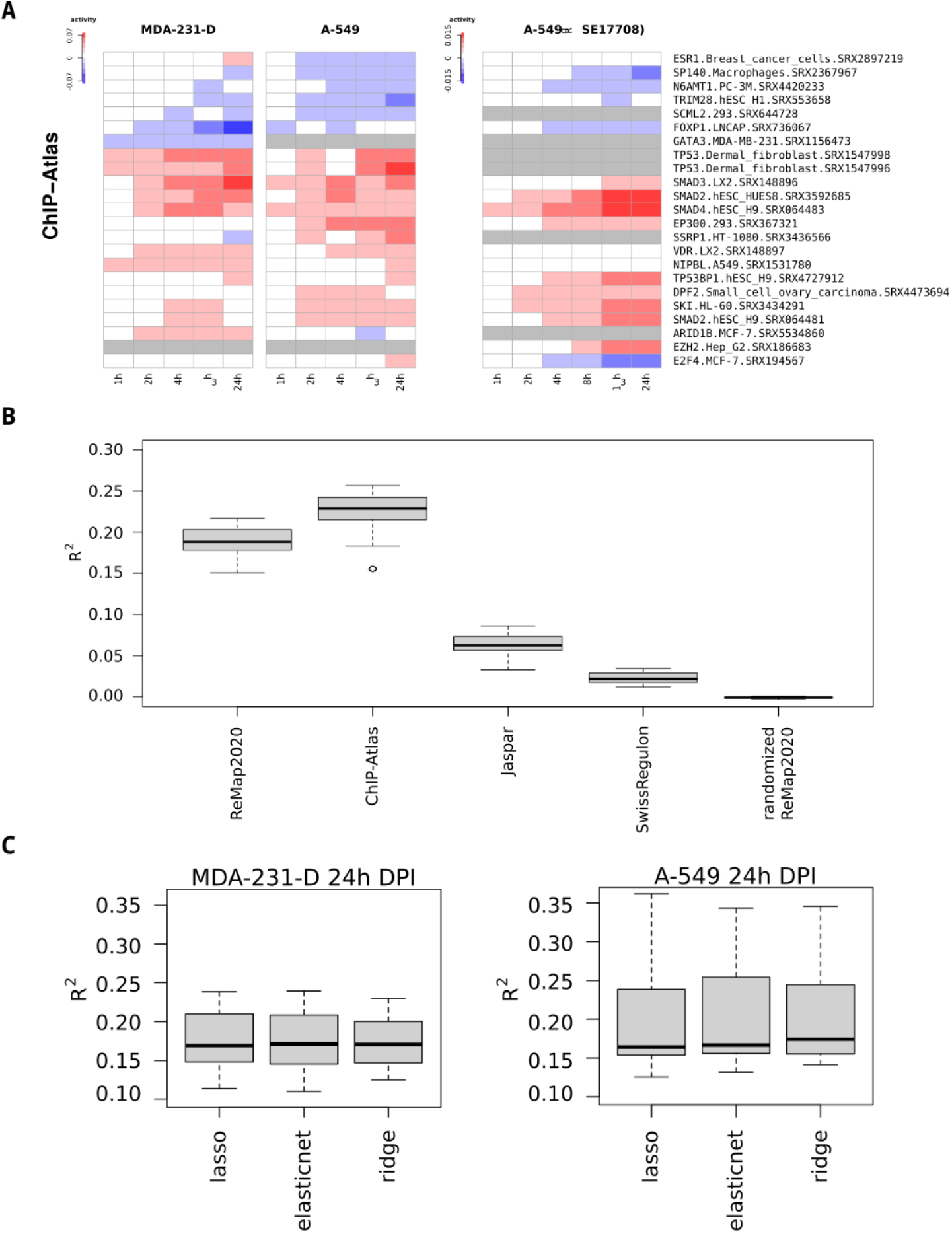
**(A)** Heatmap showing the changes of TF activities versus 0h time point during TGFβ induced EMT in A-549 and MDA-231-D cell lines. The top-scoring ChIP-Atlas are shown. Gray color designates NA values. **(B)** Comparison of models describing gene expression changes between 0 and 24 hours after rinderpest infection treatment in 293SLAM. Models were constructed using different molecular signatures sets. **(C)** Boxplots present R^2^ calculated in 10-fold cross-validation across 4 replicated experiments for models describing gene expression changes between 0 and 24 hours after TGFβ treatment in A-549 and MDA-231-D. The models were constructed using ReMap2020 molecular signatures and DPI level expression data. Lasso, elastic net and ridge regression methods were used for comparison.

**Supplementary Figure 2.**
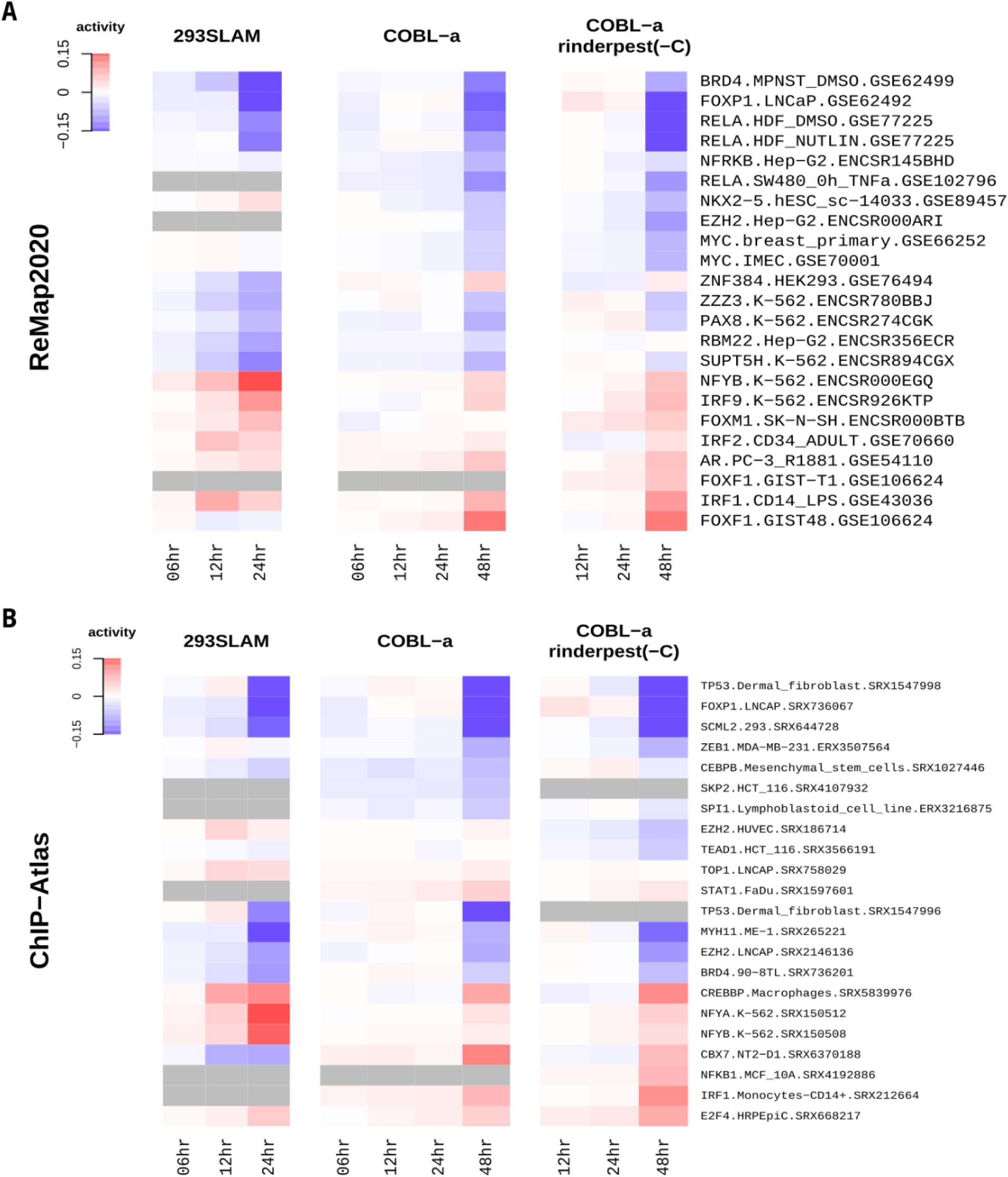
Estimating transcription factors activities from gene expression during rinderpest infection in 293SLAM and COBL-a cell lines. **(A-B)** Heatmaps present estimated replicate average activities of the most significant molecular signatures. Signature sets were constructed from ChIP-seq experiments available in ReMap2020 and ChIP-Atlas databases, respectively.

## Extended Materials and Methods

### TGF-β1 stimulation to A-549/MDA-231-D

A-549 Lung cancer cells (CCL-185, ATCC) and MDA-231-D highly metastatic human breast cancer cells (Ehata *et al*., 2007) (gift from Dr. Kohei Miyazono, Tokyo Univ.) were cultured in Dulbecco’s modified Eagle’s medium (Wako Pure Chemical Industries, Ltd, Osaka, Japan) supplemented with 10% fetal bovine serum and penicillin/streptomycin (100 U/mL, 100 µg/mL; Thermo Fisher Scientific Inc., Waltham, MA, USA). TGF-β1 (7754-BH, Recombinant Human TGF-beta 1, R&D Systems) was added at the final concentration of 1mg/mL. At 0, 1, 2, 4, 6, and 24 hours post stimulation, cells were harvested followed by RNA extraction using RNeasy mini kit (Qiagen, Valencia, CA, USA). Transcriptome data was produced by nAnT-iCAGE (Murata *et al*., 2014). CAGE libraries were sequenced on Illumina HiSeq 2500 (50-nt single read).

### Expression data processing

Raw sequencing CAGE data were processed using MOIRAI pipeline (Hasegawa et al., 2014). In short, TagDust2 was used to trim the adapter sequences and trimmed reads were then mapped to the human genome using BWA aligner. Uniquely mapping reads with overlapping 5’ ends overlapping the coordinates of FANTOM5 robust promoter set (FANTOM Consortium, 2014) were counted (raw expression counts). Counts for rinderpest infection time-course were obtained from FANTOM5’s atlas (Lizio *et al*., 2015, p. 5). For the microarray dataset we downloaded the log-transformed normalized data (GSE17708, Sartor et al. 2010).

For both CAGE datasets we consider two levels of expression data, promoter level where the expression is measured at all FANTOM5’ DPI regions (Lizio *et al*., 2015, p. 5) and gene level (HUGO gene symbol) where for each gene we use expression at its highest scored FANTOM5’ DPI region with an GENCODE 38 annotation (Frankish *et al*., 2021) and ROADMAP promoter confirmation (Kundaje *et al*., 2015). Expression tags for each sample were filtered to exclude lowly expressed promoters, normalized for the library size and transformed into counts per million (CPM) using edgeR (Robinson, McCarthy and Smyth, 2010). Next, CPM were log2 transformed with addition of pseudo count 1. For the microarray dataset we consider only the gene level expression data using already pre-normalized log-transformed data. Individual probes were matched to FANTOM5’ DPI regions based on their ENTREZID. For each dataset we designate the base level samples, taken as an earliest point in the time series, for which we calculate per gene mean expression over all replicates, further called basal expression level.

### Molecular signatures generation

We downloaded sets of peaks for all human transcription factors for human genome assembly hg38 from ReMap2020 (Chèneby *et al*., 2020, p. 20) and ChIP-Atlas (Oki *et al*., 2018) databases. For ReMap2020 no further processing of peaks was applied. In the case of ChIP-Atlas we excluded ambiguous experiments such as those labeled as “Epitope tags” or “Biotin”. The molecular signatures were constructed by first extending FANTOM5’ DPI regions by 500bp in both directions. Next, peaks were intersected with promoter regions yielding a molecular signature, where 1 indicates presence of a signature in the promoter and 0 indicates it’s absence.

Additionally, we considered molecular signatures based on predicted transcription factors binding sites from Jaspar (Fornes *et al*., 2020) and SwissRegulon (Pachkov *et al*., 2013) databases.

### Expression modeling

We describe the relationship between the expression (Y) and molecular signatures (X) using linear model formulation.

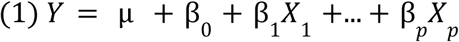

where, Y - sample expression level, *μ* - basal expression level, β_0_ - intercept, β_j_ - j-th molecular signature activity, X_j_ - j-th molecular signature.

Here, we are interested in finding unknown molecular signatures activities (β) that describe the effect of molecular signature (X) on expression (Y). By including *μ* in (1) we effectively model the change in expression between the basal expression level and corresponding sample. For each sample a separate model is fitted giving sample specific β estimates.

Models are trained using penalized linear regression. In particular, we use ridge regression (Hoerl and Kennard, 1970) which we observed to work equally well to lasso and elastic net regression (Supp. Fig. 1C). In practice, to fit our linear models we use the popular R package glmnet (Friedman, Hastie and Tibshirani, 2010).

To estimate significance of molecular activities estimates we use a test of significance for ridge regression coefficient estimates introduced by (Cule, Vineis and De Iorio, 2011), that further improves on the test proposed by (Halawa and El Bassiouni, 2000). Briefly, this test uses the normal distribution to test significance of ridge regression coefficients, assuming that under H_0_ the test statistic follows N(0, 1). For further details on standard error estimates calculation we refer interested readers to (Cule, Vineis and De Iorio, 2011), which also offers their method available as an R package. Here, we reimplement their method to facilitate combining with glmnet package.

Gene expression studies make use of experiment replication. To take advantage of these we average the activities estimates over replicates and calculate pooled standard error. We calculate pooled standard error following (Cohen, 1977).

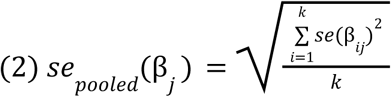

where, β_j_ - j-th molecular signature activity, se(β_ij_) - estimate of β_j_ standard error in i-th replicate, k - total number of replicates. Finally, test statistics pooled are calculated using replicate average molecular signature activities.

### Assessing Model Accuracy

To assess models accuracy we calculate R^2^ using the following formula:

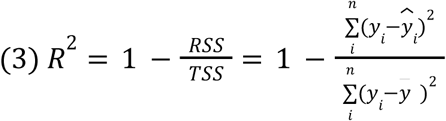

where, RSS - residual sum of squares, TSS - total sum of squares, y_i_ - i-th promoter expression, ŷ- i-th promoter predicated expression, 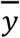 - mean promoter expression. For each model R^2^ is calculated using 10-fold cross-validation separately for each biological replicate, finally estimates are averaged across replicates.

## Notes

https://github.com/bkaczkowski/xcore

### Competing Interest Statement

The authors have declared no competing interest.

## References

Balwierz, P.J. et al. (2014) ‘ISMARA: automated modeling of genomic signals as a democracy of regulatory motifs’, Genome Research, 24(5), pp. 869–884. doi:10.1101/gr.169508.113.

Chèneby, J. et al. (2020) ‘ReMap 2020: a database of regulatory regions from an integrative analysis of Human and Arabidopsis DNA-binding sequencing experiments’, Nucleic Acids Research, 48(D1), pp. D180–D188. doi:10.1093/nar/gkz945.

Cule, E., Vineis, P. and De Iorio, M. (2011) ‘Significance testing in ridge regression for genetic data’, BMC Bioinformatics, 12(1), p. 372. doi:10.1186/1471-2105-12-372.

Dardenne, E. et al. (2014) ‘RNA helicases DDX5 and DDX17 dynamically orchestrate transcription, miRNA, and splicing programs in cell differentiation’, Cell Reports, 7(6), pp. 1900–1913. doi:10.1016/j.celrep.2014.05.010.

FANTOM Consortium et al. (2009) ‘The transcriptional network that controls growth arrest and differentiation in a human myeloid leukemia cell line’, Nature Genetics, 41(5), pp. 553–562. doi:10.1038/ng.375.

Hoerl, A.E. and Kennard, R.W. (1970) ‘Ridge Regression: Biased Estimation for Nonorthogonal Problems’, Technometrics, 12(1), pp. 55–67. doi:10.2307/1267351.

Kanehisa, M. et al. (2010) ‘KEGG for representation and analysis of molecular networks involving diseases and drugs’, Nucleic Acids Research, 38(Database issue), pp. D355–360. doi:10.1093/nar/gkp896.

Lavin, D.P. and Tiwari, V.K. (2020) ‘Unresolved Complexity in the Gene Regulatory Network Underlying EMT’, Frontiers in Oncology, 10, p. 554. doi:10.3389/fonc.2020.00554.

Lizio, M. et al. (2015) ‘Gateways to the FANTOM5 promoter level mammalian expression atlas’, Genome Biology, 16(1), p. 22. doi:10.1186/s13059-014-0560-6.

Love, M.I., Huber, W. and Anders, S. (2014) ‘Moderated estimation of fold change and dispersion for RNA-seq data with DESeq2’, Genome Biology, 15(12), p. 550. doi:10.1186/s13059-014-0550-8.

Madsen, J.G.S. et al. (2018) ‘Integrated analysis of motif activity and gene expression changes of transcription factors’, Genome Research, 28(2), pp. 243–255. doi:10.1101/gr.227231.117.

Natarajan, A. et al. (2012) ‘Predicting cell-type–specific gene expression from regions of open chromatin’, Genome Research, 22(9), pp. 1711–1722. doi:10.1101/gr.135129.111.

Oki, S. et al. (2018) ‘ChIP-Atlas: a data-mining suite powered by full integration of public ChIP-seq data’, EMBO reports, 19(12), p. e46255. doi:10.15252/embr.201846255.

Robinson, M.D., McCarthy, D.J. and Smyth, G.K. (2010) ‘edgeR: a Bioconductor package for differential expression analysis of digital gene expression data’, Bioinformatics, 26(1), pp. 139–140. doi:10.1093/bioinformatics/btp616.

Sartor, M.A. et al. (2010) ‘ConceptGen: a gene set enrichment and gene set relation mapping tool’, Bioinformatics, 26(4), pp. 456–463. doi:10.1093/bioinformatics/btp683.

Schmidt, F. et al. (2017) ‘Combining transcription factor binding affinities with open-chromatin data for accurate gene expression prediction’, Nucleic Acids Research, 45(1), pp. 54–66. doi:10.1093/nar/gkw1061.

Schmidt, F. and Schulz, M.H. (2019) ‘On the problem of confounders in modeling gene expression’, Bioinformatics, 35(4), pp. 711–719. doi:10.1093/bioinformatics/bty674.

Tian, B. et al. (2018) ‘The NFκB subunit RELA is a master transcriptional regulator of the committed epithelial-mesenchymal transition in airway epithelial cells’, The Journal of Biological Chemistry, 293(42), pp. 16528–16545. doi:10.1074/jbc.RA118.003662.

Xu, J., Lamouille, S. and Derynck, R. (2009) ‘TGF-β-induced epithelial to mesenchymal transition’, Cell research, 19(2), pp. 156–172. doi:10.1038/cr.2009.5.

## References

Cohen, J. (1977) ‘CHAPTER 2 - The t Test for Means’, in Cohen, J. (ed.) Statistical Power Analysis for the Behavioral Sciences. Academic Press, pp. 19–74. doi:10.1016/B978-0-12-179060-8.50007-4.

Ehata, S. et al. (2007) ‘Ki26894, a novel transforming growth factor-β type I receptor kinase inhibitor, inhibits in vitro invasion and in vivo bone metastasis of a human breast cancer cell line’, Cancer Science, 98(1), pp. 127–133. doi:10.1111/j.1349-7006.2006.00357.x.

Fornes, O. et al. (2020) ‘JASPAR 2020: update of the open-access database of transcription factor binding profiles’, Nucleic Acids Research, 48(D1), pp. D87–D92. doi:10.1093/nar/gkz1001.

Frankish, A. et al. (2021) ‘GENCODE 2021’, Nucleic Acids Research, 49(D1), pp. D916–D923. doi:10.1093/nar/gkaa1087.

Friedman, J.H., Hastie, T. and Tibshirani, R. (2010) ‘Regularization Paths for Generalized Linear Models via Coordinate Descent’, Journal of Statistical Software, 33, pp. 1–22. doi:10.18637/jss.v033.i01.

Halawa, A.M. and El Bassiouni, M.Y. (2000) ‘Tests of regression coefficients under ridge regression models’, Journal of Statistical Computation and Simulation, 65(1–4), pp. 341–356. doi:10.1080/00949650008812006.

Kundaje, A. et al. (2015) ‘Integrative analysis of 111 reference human epigenomes’, Nature, 518(7539), pp. 317–330. doi:10.1038/nature14248.

Murata, M. et al. (2014) ‘Detecting Expressed Genes Using CAGE’, in Miyamoto-Sato, E. et al. (eds) Transcription Factor Regulatory Networks: Methods and Protocols. New York, NY: Springer (Methods in Molecular Biology), pp. 67–85. doi:10.1007/978-1-4939-0805-9_7.

Pachkov, M. et al. (2013) ‘SwissRegulon, a database of genome-wide annotations of regulatory sites: recent updates’, Nucleic Acids Research, 41(D1), pp. D214–D220. doi:10.1093/nar/gks1145.

